# High-temporal resolution metabolic connectivity resolved by component-based noise correction

**DOI:** 10.1101/2025.08.18.670788

**Authors:** Murray B Reed, Samantha Graf, Matej Murgaš, Benjamin Eggerstorfer, Christian Milz, Leo R Silberbauer, Pia Falb, Elisa Briem, Alexandra Mayerweg, Gabriel Schlosser, Sebastian Klug, Lukas Nics, Godber M Godbersen, Sazan Rasul, Marcus Hacker, Andreas Hahn, Rupert Lanzenberger

## Abstract

Recent advances in functional PET (fPET) allow for accurate modelling of metabolic processes with a temporal resolution in the range of seconds. This enables new applications such as imaging molecular connectivity at temporal resolutions comparable to fMRI. However, high-temporal resolution fPET data are more sensitive to noise and the extraction of a meaningful signal remains a challenge.

We developed a component-based preprocessing approach adapted from fMRI, which models structured noise using tissue-specific regressors and removes low-frequency uptake trends from the fPET signal (CompCor). We applied this method to 20 high-temporal [^18^F]FDG fPET scans from a next-generation long-axial field of view PET/CT system (1s frames) and 16 scans from a conventional PET/MR scanner (3s frames). We compared filtering methods across frequency bands and examined their effects on metabolic connectivity (M-MC) estimates.

Metabolic connectivity was markedly influenced by filtering strategy and scanner type. The CompCor filter produced more consistent and structured networks than standard bandpass filters. Intermediate frequency bands (0.01-0.1 Hz) yielded the most reliable connectivity patterns between PET/CT and PET/MR data (r=0.89). High sensitivity PET/CT data revealed structured connectivity patterns also at a higher frequency band (0.1-0.2 Hz). Compared to fMRI functional connectivity, fPET-derived networks were more spatially cohesive but less differentiated.

High-temporal [^18^F]FDG fPET enables reliable estimation of individual resting-state M-MC when paired with appropriate denoising. Scanner choice and preprocessing significantly affect signal quality and interpretation, whereas the proposed physiologically informed pipeline improves comparability across systems and studies.

## Introduction

Understanding how brain regions interact remains a core question in neuroscience. Functional connectivity, measured with blood oxygenation level dependent (BOLD) functional magnetic resonance imaging (fMRI), has offered one way to study these interactions by capturing the temporal correlation of hemodynamic signals ^1,2^. However, the BOLD signal is an indirect measure of neuronal activity ^3^. It reflects a combination of changes in blood flow, blood volume and oxygen consumption rather than the specific metabolic demands that sustain neural function ^4–6^.

In contrast, [^18^F]fluorodeoxyglucose positron emission tomography ([^18^F]FDG PET) provides a more direct measure of glucose metabolism ^7^. FDG-PET has traditionally been used to create static maps of regional glucose uptake, offering specific insight into brain function. While clinically useful, static PET lacks the temporal resolution to capture moment-to-moment changes that define dynamic brain function ^8^. Recent advances have started to shift this limitation using functional PET (fPET) ^9,10^. The adoption of bolus plus constant infusion protocols, along with improved image reconstruction and processing, has increased the temporal resolution of fPET from minutes to seconds ^11,12^. With appropriate acquisition and reconstruction, metabolic signals can now be sampled in high-temporal frames, making it possible to detect acute task-evoked changes in metabolic activity ^13^.

Beyond task-based responses, this framework has sparked interest in studying resting-state metabolic connectivity (M-MC) ^14–18^. As resting-state fMRI reveals functional networks in absence of external stimulation, high-temporal resolution fPET offers a promising approach to characterize the metabolic network counterpart. This in turn could provide a molecular-level view of brain network structure in healthy individuals but also in clinical populations, complementing the hemodynamic signal and capturing aspects of neural function not seen in the fMRI signal ^19^. Previous studies have demonstrated the feasibility of estimating metabolic connectivity using [¹⁸F]FDG fPET with a temporal resolution of 16 s ^15^. However, insights from resting-state fMRI suggest that spontaneous neuronal fluctuations contributing to functional connectivity predominantly occur at higher frequencies. This raises the question of whether increasing the temporal resolution of fPET could improve the characterization of dynamic metabolic network interactions. However, such a shift toward higher temporal resolution also introduces methodological challenges, including reduced signal-to-noise ratio and increased sensitivity to preprocessing choices. Yet the shift to high temporal resolution fPET also introduces analytical challenges ^20^. Short image frames in the range of seconds substantially reduce detected counts, thereby lowering the signal-to-noise ratio and increasing vulnerability to motion, physiological artefacts, and scanner variability ^20,21^. Crucially, resting-state analyses rely on data-driven methods that are more sensitive to artifacts and the corresponding preprocessing choices ^22–26^. Thus, the ability to detect reliable connectivity patterns at rest depends heavily on how the data is filtered and denoised.

Existing PET preprocessing pipelines have been developed for lower temporal resolution ^27^. While still effective in that setting, they may not preserve the temporal structure needed for connectivity analyses at higher temporal resolution. In this case, filtering becomes a trade-off between reducing noise and preserving the signal. Too much filtering may remove biologically relevant fluctuations. Too little, and noise may falsely contribute to connectivity signals ^28,29^. The problem is further complicated by limited knowledge of which frequency bands are most relevant for metabolic signals.

The present study aims to address key analytical limitations in high-temporal-resolution [¹⁸F]FDG fPET by: (i) introducing an integrated filtering framework for robust assessment of moment-to-moment fluctuations; (ii) computing metabolic connectivity at a previously unprecedented temporal resolution of 1 s; (iii) evaluating the contribution of distinct frequency bands to metabolic connectivity; and (iv) assessing generalizability by comparing results with data from a conventional PET/MR system. These advancements are enabled by applying the proposed method to data acquired on a next-generation large axial field-of-view (LAFOV) PET/CT scanner operating in ultra-high-sensitivity mode. The integrated filtering approach, adapted from the fMRI domain, consolidates multiple preprocessing steps, including temporal detrending, regression of motion and physiological noise, and frequency-specific band separation, into a unified temporal filter. Together, our work aims to provide a robust and widely applicable approach to assess metabolic connectivity in the human brain.

## Methods

### Participants

Twenty healthy participants (mean age ± SD = 24.8 ± 3.6 years; 11 female) underwent a single [^18^F]FDG PET/CT scan using a next generation LAFOV scanner (Siemens Vision Quadra), followed by an fMRI scan on a Siemens MAGNETOM Prisma 3T. An additional 16 healthy volunteers (mean age ± SD = 26.9 ± 8.2 years; 7 female) completed two [^18^F]FDG scans on a 3T PET/MR system (Siemens Biograph mMR, Siemens Healthineers, Germany).

General health was evaluated through a structured medical assessment. This included medical history, physical examination, electrocardiogram, and routine laboratory testing. Psychiatric screening was conducted using the Structured Clinical Interview for DSM-IV Axis I Disorders (SCID-I) to rule out current or past psychiatric diagnoses.

Exclusion criteria included any current or previous severe medical condition, psychiatric disorder, psychopharmacological treatment (past 6 months), or contraindications to PET/MRI. These included metallic implants, claustrophobia, and prior radiation exposure. Urine drug tests were performed at screening. For female participants, urine pregnancy tests were conducted at screening and again before each scan.

All participants provided written informed consent and received financial compensation for their time. Both studies were approved by the ethics committee of the Medical University of Vienna (EK 1307/2014 and 1642/2022) and conducted in accordance with the Declaration of Helsinki. The investigation forms part of a larger study registered prior to participant recruitment at clinicaltrials.gov (NCT02711215 and NCT06243783, respectively).

### Study Design

For the LAFOV PET/CT study, participants were positioned with the head centered in the scanner bore to optimize count sensitivity. Data was acquired in ultra-high sensitivity mode (∼176 kcps/MBq ^30,31^). Each session began with a topogram and a low-dose CT scan, followed by a 25-minute resting-state PET acquisition ^32^. During the scan, participants were instructed to remain awake with their eyes open, maintain fixation on a black crosshair presented on a gray background, and allow their thoughts to wander freely without focusing on any specific task. Radiotracer administration followed a bolus plus constant infusion protocol, initiated simultaneously with the start of PET acquisition (see data acquisition).

In the separate PET/MR cohort, participants underwent two sessions as part of a randomized, double-blind, cross-over design involving administration of either the selective serotonin reuptake inhibitor citalopram or a placebo on a Siemens Biograph mMR 3T (∼15.0 kcps/MBq ^33^). Each session began with a high-resolution T1-weighted structural MRI scan. This was followed by a 50-minute simultaneous PET and BOLD-fMRI acquisition. The radiotracer protocol was identical to the PET/CT study, with bolus and infusion starting at the beginning of the scan. At 20 minutes, the pharmacological agent was administered intravenously. Throughout the acquisition, participants maintained visual fixation and were instructed to remain still and let their thoughts wander. For the present analysis, only the first 20 minutes of the fPET data from the placebo session were included. This matched the acquisition window of the PET/CT study and ensured that pharmacological effects did not confound the results.

### Data acquisition and reconstruction

[^18^F]FDG was synthesized on each study day at the Department of Biomedical Imaging and Image-guided Therapy, Division of Nuclear Medicine, Medical University of Vienna.

At the start of fPET acquisition, [^18^F]FDG was injected through a cubital vein as a 1-minute bolus, followed by constant infusion for the rest of the scan using a shielded pump (Syramed mSP6000, Arcomed, Switzerland). The dose was 5.1 MBq per kg body weight. Bolus speed was 816 ml per hour. Infusion speed was 91.0 ml per hour for the LAFOV system and 29.5 ml per hour for the PET/MR system, with a bolus and infusion ratio of 20:80 ^12^. All fPET data on the PET/MR system were acquired in list mode, allowing frame lengths to be defined during reconstruction.

The T1-weighted structural image acquired utilized a magnetization-prepared rapid gradient echo (MPRAGE) sequence (TE 4.21 ms, TR 2200 ms, voxel size 1 x 1 x 1.1 mm, matrix 240 x 256, 160 slices, flip angle 9°, TI 900 ms, duration 7.72 minutes).

In the LAFOV PET/CT protocol, a low-dose CT was acquired after breath-hold instructions (120 kVp, 20 mA, CareDose4D and CarekV enabled). CT images were reconstructed with 0.98 x 0.98 x 4 mm voxels. Thereafter, fPET data was acquired in list mode. MRI data for the LAFOV study was recorded on a Siemens Prisma 3T scanner with a 64-channel head coil. Functional MRI images for the LAFOV study were acquired with an EPI sequence (TE 30 ms, TR 900 ms, field of view 200 x 200 mm, resolution 80 x 80 pixels, 48 axial slices of 2.4 mm thickness, multiband factor 4, TA: 20:07 min).

All fPET data were reconstructed using Siemens’ E7 Tools. LAFOV fPET data was reconstructed using 3D-TOF OP-OSEM with 4 iterations and 5 subsets into 1500 frames of 1s. CT-based attenuation correction was applied. PET/MR data was reconstructed with OP-OSEM using 3 iterations and 21 subsets into 400 frames of 3s each. Here, attenuation correction was based on a pseudo-CT derived from the T1 image ^34^.

For the comparison between LAFOV PET/CT and PET/MR data, the LAFOV data (1s frames) was downsampled after reconstruction to match the 3s frame duration of the PET/MR acquisitions.

### CompCor Filter

To remove structured noise from the fPET data and improve signal quality, we implemented a variant of the data-driven CompCor (component-based noise correction) method, originally developed for fMRI ^23^, and adapted it for use with fPET data. This approach is designed to reduce physiological and motion-related noise by identifying and removing components of the signal that are unlikely to originate from neural activity. Specifically, fluctuations associated with white matter, cerebrospinal fluid (CSF), and head motion were treated as nuisance regressors. Motion effects were modeled using the Friston 24-parameter model, which includes the six rigid-body motion parameters, their temporal derivatives, and the corresponding squared terms ^35^. Tissue-specific nuisance signals were extracted from SPM’s tissue probability maps, with thresholding applied to ensure exclusion of gray matter regions and minimize the inclusion of signal from tissue of interest. Principal component analysis was then used to capture the dominant patterns of non-neural variability, which were regressed out of the fPET data (see below). This also removes the cumulative tracer uptake from the signal to allow for accurate M-MC estimation ^8^. Additionally, temporal filtering was used to suppress slow drifts and high-frequency noise. This method enhances the sensitivity of molecular connectivity (MC) analyses by reducing structured noise and isolating signal changes more closely associated with underlying neural processes. See Figure 1 for a graphical overview of the CompCor method.

**Figure 1:**
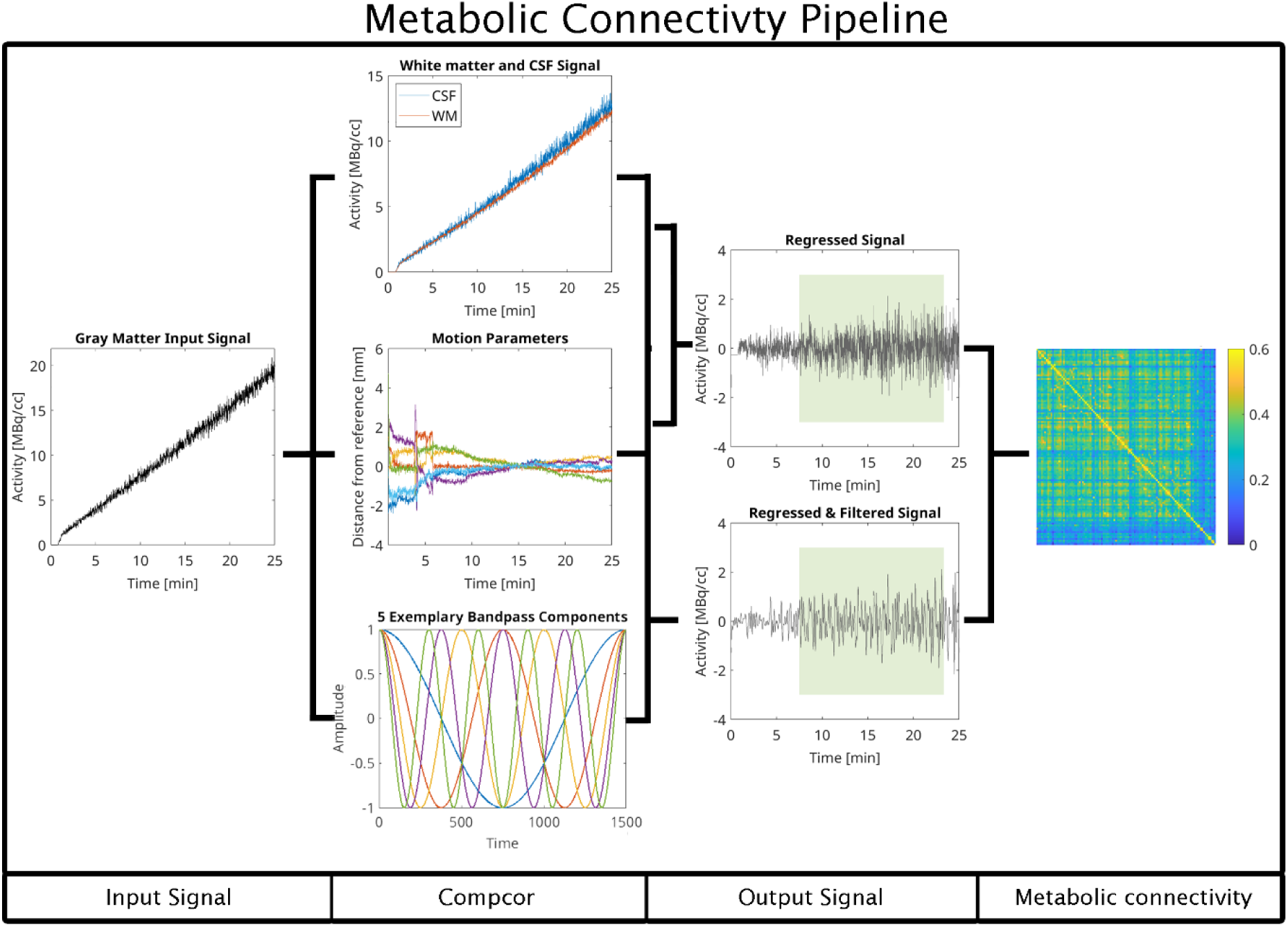
Computation of metabolic connectivity using the CompCor-based preprocessing pipeline. The approach operates in a voxelwise fashion to prepare high-temporal resolution fPET data for connectivity analysis. After standard preprocessing, the gray matter signal is isolated and submitted to the CompCor algorithm alongside head motion parameters and, optionally, a specified frequency band of interest. The algorithm first extracts nuisance signals from anatomically defined white matter and cerebrospinal fluid masks. It then derives a set of principal components that capture the majority of variance within these non-neuronal compartments, reflecting physiological and scanner-related noise. To improve correction for head motion, the six rigid-body motion parameters are expanded by computing their temporal derivatives, squared terms, and squared derivatives, again followed by principal component analysis. This extended motion model allows for the capture of both linear and nonlinear movement-related artifacts. If a bandpass filter is defined, the corresponding sine and cosine basis functions are generated to model frequencies outside the desired spectral window. All nuisance regressors, including white matter and CSF components, expanded motion parameters, and optional frequency regressors, are entered into a single multiple regression model. This simultaneous regression removes confounding sources of variance in a unified step, minimizing the risk of reintroducing noise through sequential filtering. The resulting time series are thus cleaned of motion artifacts, physiological noise, non-neuronal fluctuations, and undesired frequency components. These denoised signals provide a robust basis for the estimation of metabolic connectivity metrics.

### Preprocessing of fPET data

Preprocessing and quantification of all fPET data followed procedures established in previous studies ^12,36^. Head motion correction was performed using SPM12 (https://www.fil.ion.ucl.ac.uk/spm) with the quality setting at “best.” Images were realigned to a motion-free reference created from a stable time segment late in the scan. The mean PET image was coregistered to each participant’s structural MRI, which in turn was spatially normalized to MNI space. These transformations were then applied to the full dynamic fPET series. Normalized data were either smoothed with an 8 mm FWHM Gaussian kernel or filtered with an extended dynamic non-local means (edNLM) filter (for PET/MR data only, due to the lower scanner sensitivity) as the latter proved most efficient for task-based fPET analyses ^37^. The edNLM filter used kernels of 3x3x3 voxels × 3 frames and 3x3x3 voxels × 5 frames, followed by 5 mm Gaussian smoothing ^37^.

After preprocessing, the time series from each participant was filtered voxel-wise using the CompCor method, or region-wise using a second-order Butterworth band-pass filter. The Butterworth filter was implemented as an infinite impulse response filter in Matlab and included correction for motion parameters through partial correlation. The data were filtered into the following frequency bands (see results): 0.001 - 0.01 Hz, 0.01 - 0.1 Hz, and 0.1 - 0.2 Hz for the LAFOV dataset, and 0.001 - 0.01 Hz, 0.01 - 0.1 Hz, and 0.1 - 0.16 Hz for the PET/MR dataset.

Band limits for the PET/MR data were adjusted to remain within the Nyquist frequency. These bands were selected by examining the power spectral density of each scan to identify the frequency ranges where signal peaks were observed.

Metabolic connectivity matrices were then calculated from the processed time series. For each participant, regional mean signals were extracted using a combined parcellation of the Schaefer 100 cortical parcels ^38^ and 14 subcortical regions from the Harvard-Oxford atlas ^39^. Pearson correlation was used to compute the M-MC matrix for each subject filtered by CompCor. For Butterworth-filtered data, partial correlation was applied to account for motion parameters, which is inherently processed in the CompCor.

### Preprocessing of fMRI data

Physiological noise in the fMRI data (acquired at Siemens Prisma scanner) was reduced using PESTICA ^40^. Further preprocessing was performed in SPM12. Slice-timing correction was applied relative to the temporally middle slice. Images were then realigned to the mean image.

Spatial normalization used the standard MNI template. Images were resliced to 2.5 mm isotropic resolution, preserving approximate voxel volume. Nonlinear artifact reduction was carried out with the BrainWavelet Toolbox ^41^, using the “chsearch” parameter set to “harsh” and a threshold of 20. This was optimized for unsmoothed data acquired with multi-band acceleration, which typically has reduced signal-to-noise ratio. All images were masked using a gray-matter template. Smoothing was applied using a Gaussian kernel three times the resliced voxel size (8mm).

Following preprocessing, CompCor filtering was applied in a voxel-wise manner, similar to fPET. Regional mean signals were extracted using the same atlas parcellation as for fPET (Schaefer 100 + Harvard-Oxford subcortical). Functional connectivity was estimated using Pearson correlation on the data after removing the initial 5min of data points.

### Statistical analysis

Mean metabolic and functional connectivity matrices were computed across subjects for visualization purposes. To assess the similarity between filtering approaches, Spearman correlations were calculated between CompCor and Butterworth filtering, and between Gaussian smoothing and edge-preserving non-local means (edNLM) filtering. Additional comparisons were made between scanner types (PET/MR vs LAFOV) and across frequency bands.

To investigate macroscale network structure of fPET and fMRI data, connectivity matrices were subjected to hierarchical clustering. The clustering used Ward’s linkage method and Euclidean distance as the similarity metric. The quality of the resulting dendrograms was evaluated using the cophenetic correlation coefficient, which quantifies how faithfully the clustering preserves the pairwise distances between elements.

In addition, connectivity matrices were clustered using agglomerative clustering with eight predefined clusters. This choice was based on the Yeo 7-network cortical parcellation ^42^, with one additional cluster assigned to subcortical regions. The goal was to enable a more direct comparison between metabolic and functional network structures.

## Results

High-temporal resolution metabolic fPET data acquired with the next-generation LAFOV PET/CT scanner exhibited distinct spectral features following baseline uptake correction. Power spectral density (PSD) analyses revealed individual and group-level peaks across multiple low-frequency bands (Figure 2a and supplementary figure S1).

**Figure 2:**
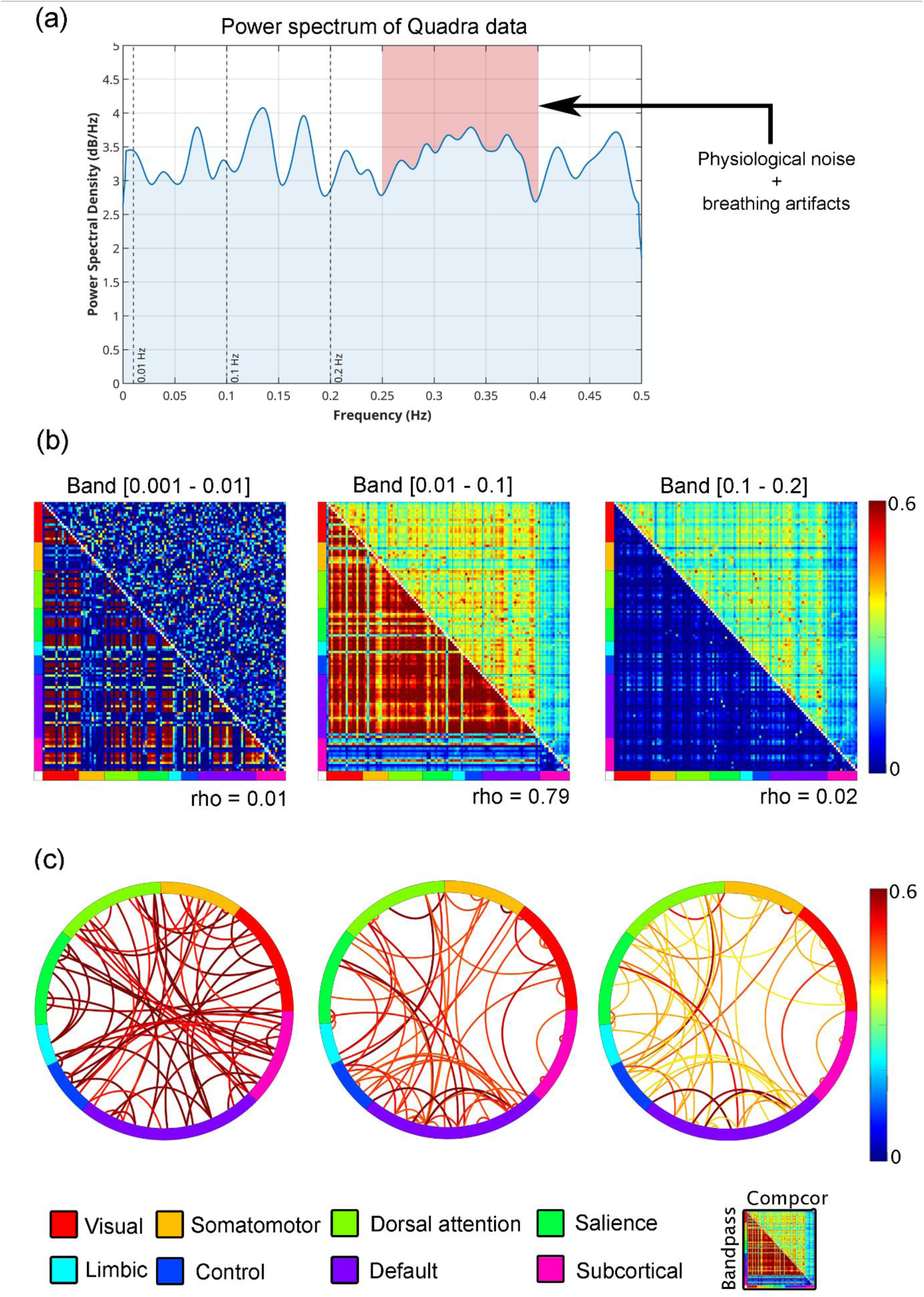
Overview of metabolic connectivity metrics estimated using CompCor and Butterworth band-pass filtering on high-temporal resolution fPET data acquired with a LAFOV PET/CT system. (a) shows an individual power spectral density estimates after removal of baseline uptake (blue). Standard frequency bands are depicted as dashed vertical lines for 0.01, 0.1 and 0.2 Hz. The red shaded area indicates the typical bands where physioligical noise such as breathing artifacts occur ^50,51^. (b) displays the metabolic connectivity matrices, separated by frequency band. The lower triangle shows connectivity derived from the Butterworth filter. The upper triangle shows connectivity estimated using the CompCor method. Connectivity estimates derived using the Butterworth filter were overestimated in the lower frequency bands (0.001 - 0.01 Hz and 0.01 - 0.1 Hz), and substantially underestimated in the higher frequency band (0.1 - 0.2 Hz). Notably, the 0.01 - 0.1 Hz band was the only range in which a strong linear correlation was observed between the two filtering methods. (c) illustrates the strongest 5 percent of connections across each frequency band, based on CompCor- filtered data. Connections are grouped according to large-scale functional networks as defined by ^52^, along with subcortical regions from the Harvard-Oxford atlas ^53^.

Comparison of filtering methods revealed notable differences in estimated M-MC (Figure 2b). While the 0.01 - 0.1 Hz band yielded high similarity between the CompCor and Butterworth filter (Spearman’s rho = 0.79), both lower (0.001 - 0.01 Hz) and higher (0.1 - 0.2 Hz) bands showed no agreement (rho ≤ 0.02). Butterworth-filtered data tended to overestimate connectivity in the lower two bands and underestimate it in the higher band relative to CompCor-filtered data. Visualization of the strongest 5% of CompCor-filtered connections highlighted distinct spatial organization across bands, consistent with known functional network architecture (Figure 2c). Notably, the 0.1 - 0.2 Hz band exhibited similarly structured resting-state-like connectivity patterns as compared to the 0.01 - 0.1 Hz band, and correlated highly (rho = 0.94) when using CompCor. For the Butterworth-filtered data, the correlation between these frequency bands was markedly lower (rho = 0.29). However, the connectivity strength was decreased as within and between network connectivity such as the default mode network was less pronounced. This suggests that meaningful metabolic organization may be most prominent within intermediate frequency ranges (e.g., 0.01 - 0.1 Hz). In contrast, the 0.001 - 0.01 Hz band showed a random-like pattern.

To examine the impact of scanner sensitivity on M-MC using the CompCor method, we compared results from the next-generation LAFOV PET/CT system with those from a standard Siemens PET/MR hybrid scanner (Figure 3). Both, 0.01 - 0.1 Hz and 0.1 - 0.16 Hz bands yielded high inter-scanner correlations (rho = 0.89 and rho = 0.86, respectively). Nonetheless, connectivity strength was consistently greater in data acquired with the next-generation LAFOV PET/CT system, indicating enhanced sensitivity to spontaneous metabolic fluctuations. Generally, the observed connectivity amplitudes are consistent with previously reported connectivity estimates from standard PET systems ^11,15^ and underscore the improved sensitivity of the LAFOV PET/CT scanner. In the 0.001 - 0.01 Hz band, scanner-derived connectivity metrics showed limited correspondence (rho = 0.28). Of note, PET/MR data exhibited the highest connectivity in this band, which however seems to be attributed to the lower sensitivity and lower sampling rate (see discussion).

**Figure 3:**
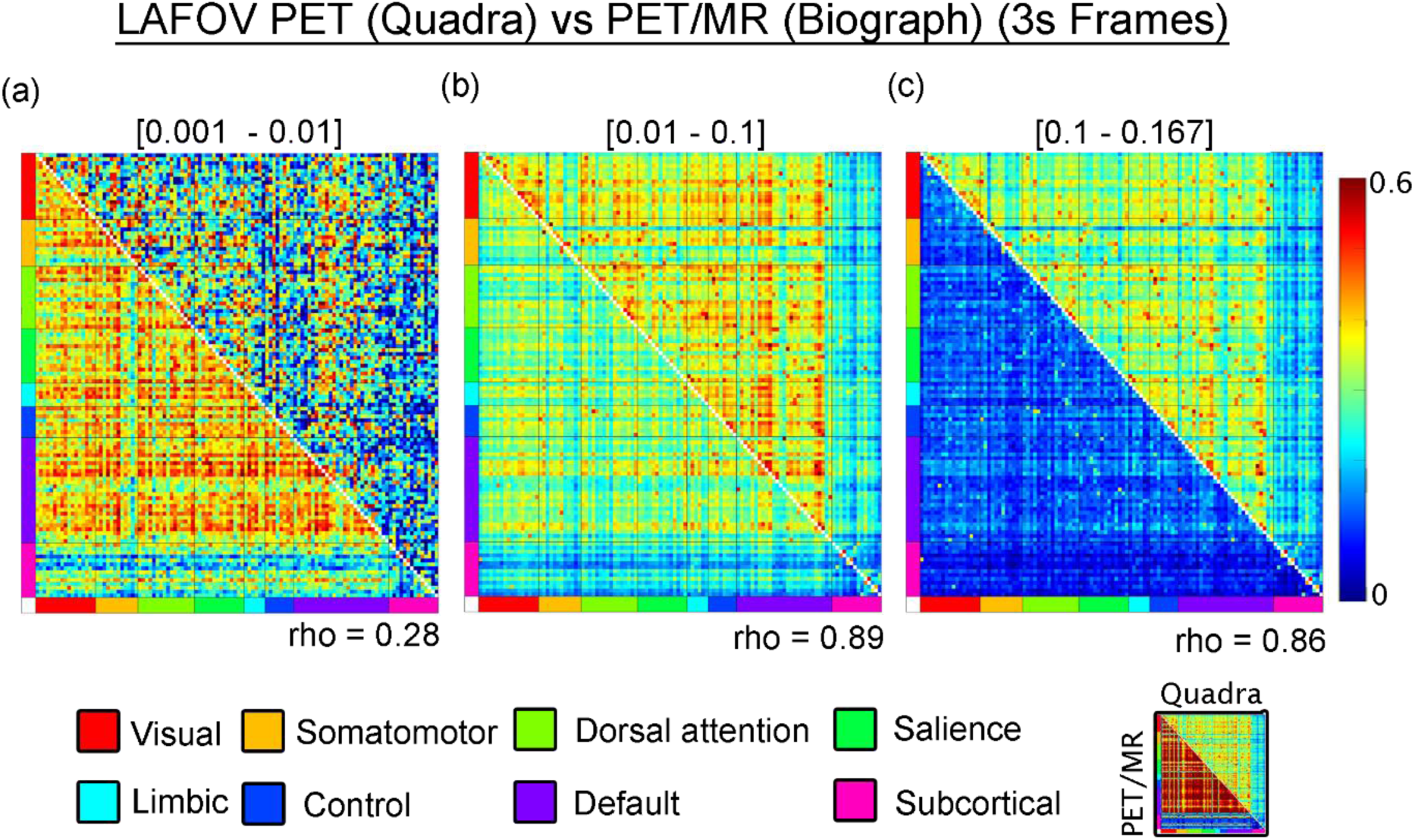
Comparison of metabolic connectivity metrics derived from a conventional PET/MR system (lower triangle) and a next-generation LAFOV PET/CT system (upper triangle) using the CompCor method. (a) Connectivity estimates in the lower frequency band (0.001 - 0.01 Hz) show a weak linear relationship between scanners (Spearman’s rho = 0.28), indicating poor correspondence in this range. (b) In the mid-frequency band (0.01 - 0.1 Hz), however, a strong linear correlation is observed between the two systems (rho = 0.89), suggesting improved consistency across scanner types. (c) The higher band (0.1 - 0.16 Hz) also shows a high correlation between systems (rho = 0.86), although connectivity estimates from the PET/MR system are generally weaker in magnitude.The lower strength of connectivity observed in the PET/MR data aligns with findings from previous studies using standard PET systems (e.g., ^11,15^). Connectivity matrices are organized according to large-scale functional networks described by ^52^, along with subcortical regions from the Harvard-Oxford atlas ^53^.

Next, the influence of spatial filtering methods was assessed. In the lowest frequency band (0.001 - 0.01 Hz), edNLM and Gaussian-smoothed data showed a moderate correlation (rho = 0.72), with comparable connectivity magnitudes. For the 0.01 - 0.1 Hz band, edNLM filtering yielded highly elevated connectivity values, yet a strong linear relationship was preserved between methods (rho = 0.91). In the highest frequency band (0.1 - 0.2 Hz), M-MC was again more pronounced in data processed with the edNLM filter. Although the spatial pattern of connectivity remained moderately consistent with Gaussian smoothing (rho = 0.72), the overestimation introduced by edNLM rendered the resulting M-MC values less comparable to those reported in previous literature, see Figure 4. A similar pattern was observed when the temporal window of the edNLM filter was reduced (Supplementary Figure 2), further indicating that the degree of temporal smoothing applied prior to M-MC estimation exerts a substantial influence on the connectivity strength.

**Figure 4:**
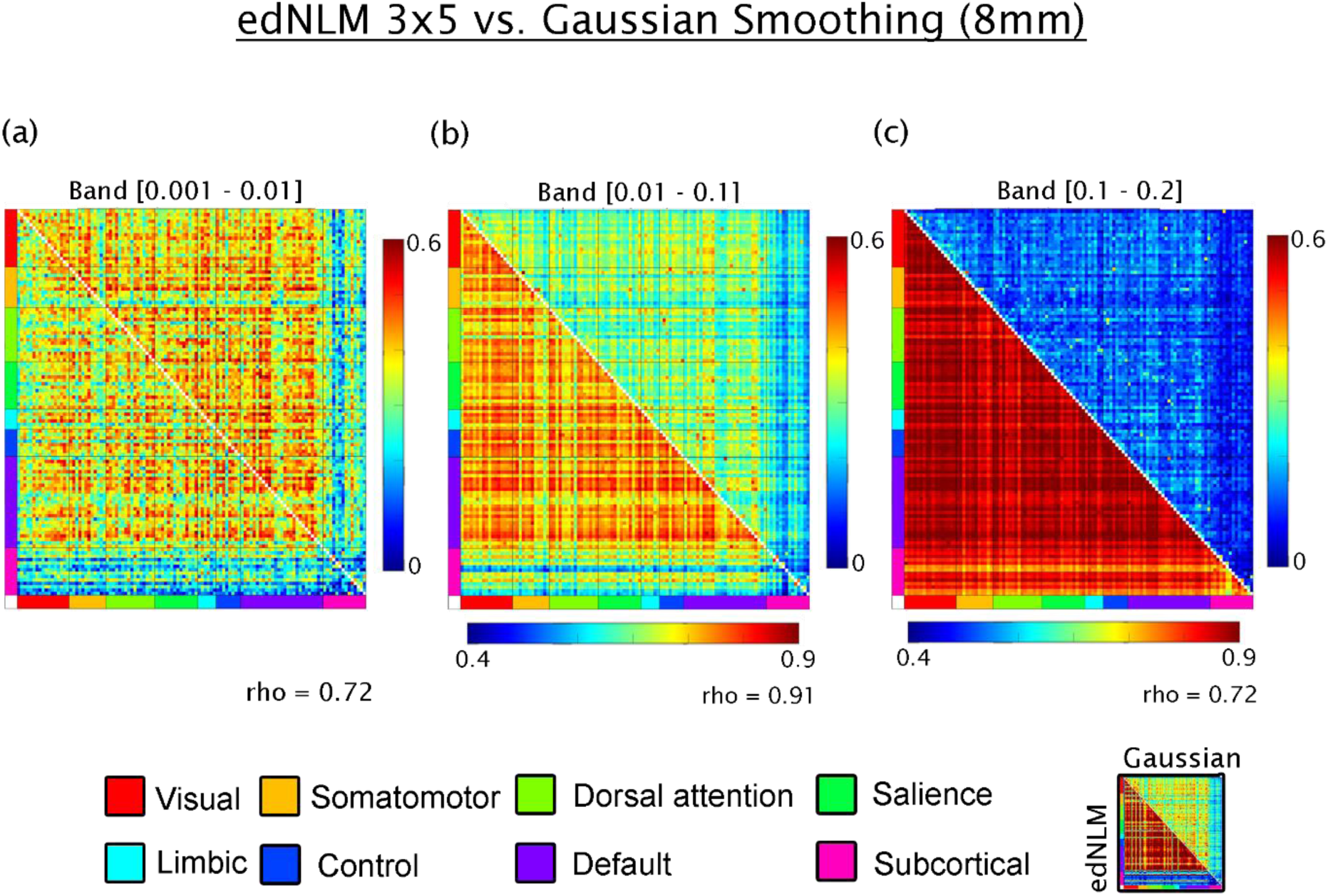
Comparison of filtering techniques applied to high-temporal resolution metabolic connectivity data acquired with a current-generation PET/MRI scanner. (a) In the lowest frequency band (0.001 - 0.01 Hz), metabolic connectivity estimates processed with extended dynamic non-local means (edNLM) filtering (upper triangle) show comparable magnitude to those processed with conventional Gaussian smoothing (lower triangle), with a moderate linear correlation between methods (Spearman’s rho = 0.72). (b) In the 0.01 - 0.1 Hz band, edNLM filtering yields noticeably higher connectivity values compared to Gaussian smoothing, although the correspondence between methods remains high (rho = 0.91). (c) In the higher frequency band (0.1 - 0.2 Hz), the linear association remains moderate (rho = 0.72), but the edNLM-processed data consistently overestimate connectivity relative to the Gaussian approach. Connectivity values are grouped according to large-scale functional networks defined by ^52^, along with subcortical regions from the Harvard-Oxford atlas ^53^.

Hierarchical clustering revealed differing organizational properties of metabolic and hemodynamic connectivity networks (Figure 5). Dendrograms constructed from fPET and fMRI connectivity matrices showed that while both modalities exhibited distinct network boundaries, M-MC clusters appeared more spatially cohesive and less dispersed. The cophenetic correlation coefficients of the resulting hierarchical trees were comparable (fPET: c = 0.60; fMRI: c = 0.62), indicating similar internal consistency.

**Figure 5:**
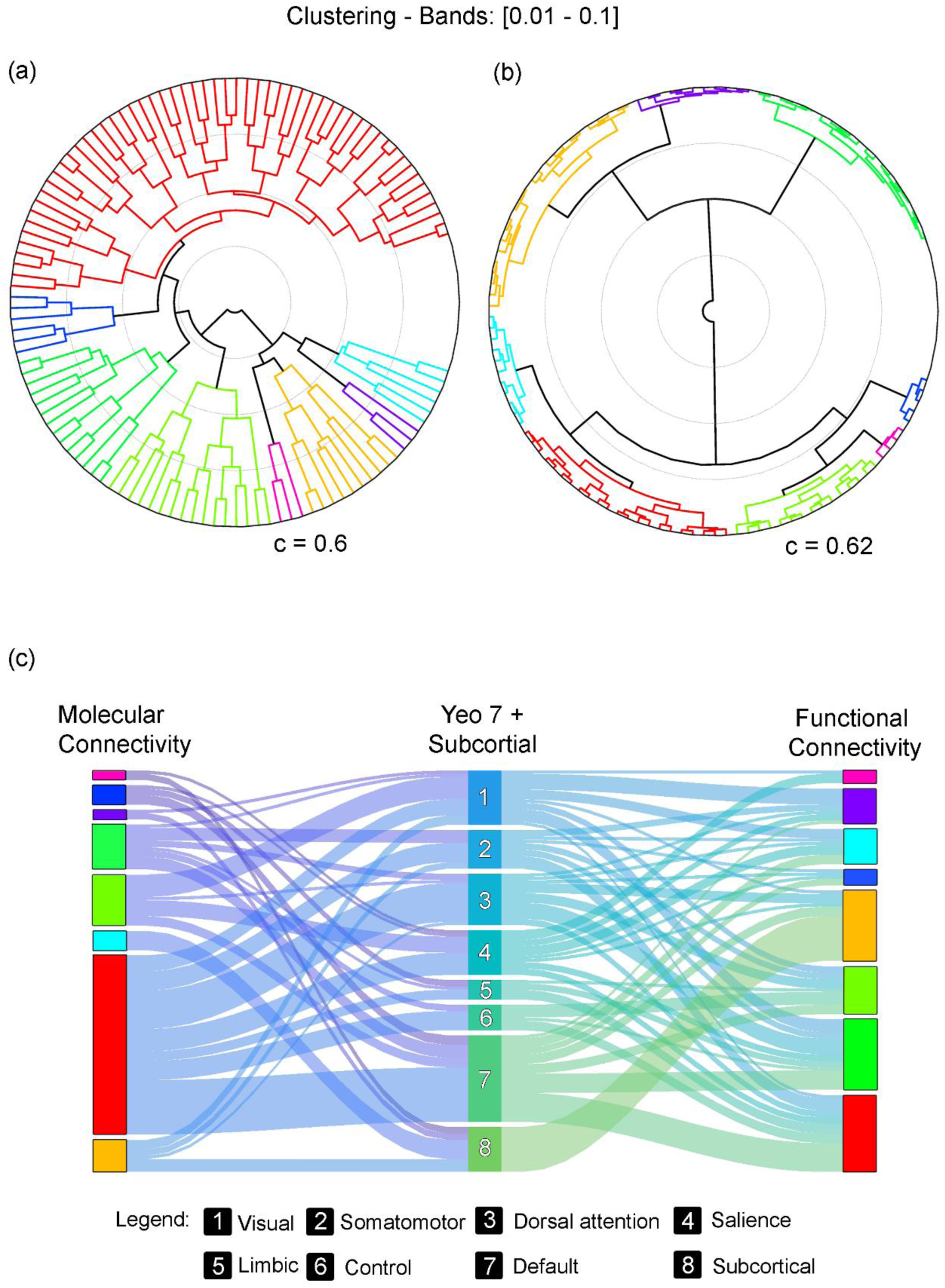
Comparison of metabolic and functional connectivity network clustering. (a) Hierarchical clustering of metabolic connectivity data acquired from the next-generation PET/CT scanner reveals distinct network structures. (b) A corresponding dendrogram illustrates the functional connectivity networks derived from fMRI data. While both modalities show clear modular organization, metabolic connectivity clusters are positioned more closely together, suggesting higher inter-network similarity compared to the more differentiated structure observed in functional connectivity. The cophenetic correlation coefficients were similar across modalities, with c = 0.60 for metabolic connectivity and c = 0.62 for functional connectivity, indicating comparable consistency in the hierarchical cluster solutions. (c) A Sankey plot visualizes the correspondence between clustered metabolic networks and canonical functional networks as defined by Yeo et al ^52^. Notably, one prominent metabolic cluster (colored red) spans across all Yeo-defined networks, suggesting a degree of global integration in the metabolic architecture not observed in the functional domain.

To facilitate direct comparison, both datasets were further partitioned into 8 clusters using agglomerative clustering, reflecting 7 canonical Yeo networks and 1 subcortical cluster. A Sankey diagram revealed partial overlap between the metabolic and functional clusters (Figure 5c), with the largest metabolic cluster spanning across all major Yeo networks, suggesting broader cross-network metabolic integration relative to fMRI.

## Discussion

This study provides a robust basis for the investigation of individual-level metabolic connectivity using high-temporal resolution [^18^F]FDG fPET data. We systematically investigated how scanner sensitivity (next-generation LAFOV PET/CT vs. conventional PET/MR), temporal resolution (1s vs. 3s), signal processing approaches (CompCor vs. standard bandpass filtering), and spatio-temporal smoothing methods (Gaussian vs. edNLM) affect the estimation of M-MC. Our results highlight key insights: (i) M-MC estimates are sensitive to both scanner performance and temporal preprocessing; (ii) the 0.01 – 0.1 Hz frequency band includes meaningful network patterns for both scanner systems, (iii) higher sensitivity of next-generation PET systems captures higher-frequency connectivity patterns that are aliased or lost in conventional PET/MR data; and (iv) applying advanced smoothing filters (e.g., edNLM) increases signal amplitude but raises concerns regarding comparability with existing literature. Despite the differences in acquisition and preprocessing, we found a high correlation between M-MC matrices across scanners in the middle frequency band, although absolute values diverged systematically.

A central methodological question addressed in this study concerns the preprocessing strategy used to isolate relevant metabolic fluctuations. Standard bandpass filtering lacks the ability to remove structured noise from physiological or scanner-related sources ^23,43,44^. In contrast, the CompCor method, which was originally developed for fMRI ^23^, offers key advantages for fPET by modeling nuisance signals from white matter and cerebrospinal fluid and regressing them from the data parallel to an optional bandpass frequency filtering. Importantly, this approach also suppresses the irreversible tracer uptake component of the FDG signal, which otherwise dominates the low-frequency spectrum without the need to choose a specific frequency band. Furthermore, our results demonstrate that the CompCor method consistently resulted in lower amplitude but more accurate M-MC estimates for PET/MR data across all frequency bands when compared to bandpass filtering. However, this suppression likely reflects a more conservative and specific isolation of dynamic fluctuations in glucose metabolism, supporting its suitability for fPET applications, particularly when combined with a temporal resolution of 3 s.

Temporal resolution emerged as a decisive factor in capturing meaningful M-MC. The LAFOV PET/CT system’s 1s framing enabled robust detection of dynamic fluctuations also in the 0.01 - 0.2 Hz range. In contrast, PET/MR data acquired at 3s resolution were inherently limited by a lower Nyquist frequency (∼0.167 Hz), resulting in aliasing of higher-frequency components. These aliased signals can misleadingly appear in low-frequency bands, as demonstrated by observed peaks around 0.033 Hz in the PET/MR data (Fig. 3a vs. c, lower triangles). Theoretical analysis confirmed that high-frequency physiological or vascular signals (e.g., respiratory signals around 0.3 Hz) can alias into this lower range due to undersampling. For a sampling interval of 3 s (PET/MR), the Nyquist frequency is 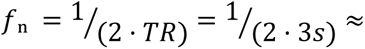. Frequencies above this threshold are folded back into lower bands according to the aliasing formula 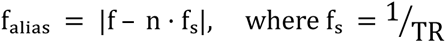 is the sampling rate. As a result, aliased components appear within the 0.001 - 0.01 Hz range and contaminate the lower bands ^45^. This presents an interpretability issue, as aliased frequencies may mimic low-frequency neural connectivity patterns, including those traditionally attributed to resting-state networks. Consequently, accurate characterization of M-MC, especially in higher frequency bands, requires high temporal resolution to avoid spectral distortion.

While the application of temporally-aware smoothing filters (e.g., edNLM) have been shown to improve fPET outcome parameters for task-specific analyses ^11^, our findings caution against the use of temporal smoothing methods in high-temporal M-MC research. Although edNLM- filtered data showed strong correlations with Gaussian-smoothed data across bands (e.g., rho = 0.72 - 0.91), the absolute values of M-MC became significantly inflated, particularly in the higher bands (0.1 - 0.2 Hz). This inflation renders direct comparisons with previously published PET connectivity metrics difficult and raises questions about the biological interpretability of such enhanced signals. Similar effects were seen when reducing the temporal component of the edNLM filter, underscoring the strong, potentially detrimental, influence of temporal smoothing on connectivity outcomes. Thus, while advanced filters may be valuable for enhancing spatial consistency, they must be calibrated carefully to preserve physiological interpretability and cross-study comparability in M-MC.

Interestingly, M-MC components, particularly those within the 0.01 - 0.1 Hz band, were still accurately detectable in PET/MR data despite the lower frame rate and scanner sensitivity ^11,15^. This reflects the robustness of certain connectivity patterns. However, the lower sampling rate (3s) and sensitivity made further spectral decomposition of other frequency bands unreliable. This finding underscores a subtle yet important point that while the spatial patterns of M-MC may persist at lower temporal resolutions, accurate frequency-resolved analysis depends critically on adequate sampling.

Furthermore, discrepancies in connectivity observed in the 0.001 - 0.01 Hz band in PET/MR data may be attributed to aliasing of higher-frequency signals. As previously discussed, respiratory or vascular dynamics in the 0.2 - 0.4 Hz range ^46^, well above the Nyquist threshold of PET/MR data, may alias into the 0.001 - 0.01 Hz range, falsely suggesting low-frequency connectivity. The data acquired using the next-generation scanner, with its higher frame rate of 1s, was able to avoid such artifacts and allowed these signals to be more accurately assigned to their true spectral domain. Thus, spectral peaks < 0.01 Hz in the PET/MR data should be interpreted cautiously, as they may represent aliased higher-frequency physiological processes rather than genuine resting-state metabolic activity.

A notable observation was that connectivity matrices derived from fPET and fMRI showed a similar modularity, both resolving canonical functional networks, but differed markedly in spatial organization. Specifically, M-MC clusters derived from fPET were more spatially cohesive and less dispersed than those from fMRI. This pattern likely reflects fundamental differences in the underlying signals: while fMRI captures rapid hemodynamic fluctuations linked to neuronal activity, fPET reflects fluctuations in regional glucose metabolism, which may be governed by integrative processes such as synaptic maintenance, glial-neuronal interactions, or neuromodulatory tone ^19,47–49^. We speculate that these ubiquitous metabolic processes may result in broader and more spatially homogeneous connectivity patterns, however, the actual underlying mechanisms still need to be assessed in future work. Nevertheless, the cophenetic correlation coefficients for both modalities were comparable (c ∼ 0.6), suggesting that both hierarchies are internally consistent, albeit representing different neurobiological processes.

Several limitations warrant consideration. First, although our results emphasize the superior temporal resolution of the LAFOV scanner, these benefits are only realized when combined with appropriate preprocessing and filtering strategies. Second, while we used CompCor and bandpass filtering as two representative approaches, the space of denoising methods for fPET is expanding and may benefit from hybrid or adaptive techniques in the future. Third, absolute connectivity values depend on normalization, smoothing, and segmentation procedures, which may vary. Lastly, the interpretation of M-MC, while increasingly supported by emerging literature ^8,19^, remains less established than in fMRI, necessitating further work to identify the underlying neurophysiological processes. Nevertheless, our work demonstrates that metabolic connectivity from high-temporal resolution fPET data can be robustly obtained from ultra-high-sensitivity LAFOV and conventional scanner systems.

In sum, we provide a comprehensive comparison of M-MC estimation across PET scanner generations and preprocessing pipelines. Our findings demonstrate that next-generation PET systems with high temporal resolution substantially improve the reliability of M-MC estimation, particularly in higher frequency bands. However, methodological choices particularly regarding filtering and denoising profoundly influence the amplitude, spatial organization, and interpretability of connectivity metrics. Our results support the use of high-temporal-resolution fPET, combined with physiologically-informed preprocessing to enhance the reliability and comparability of M-MC estimates. Altogether, this opens a new avenue in molecular imaging, enabling the assessment of metabolic connectivity at rest and during cognitive, interventional or pharmacological stimulation as well as the re-organization of metabolic network interactions in various brain disorders.

## Statements and Declarations

### Clinical trial number

NCT02711215 and NCT06243783

### Author Contribution

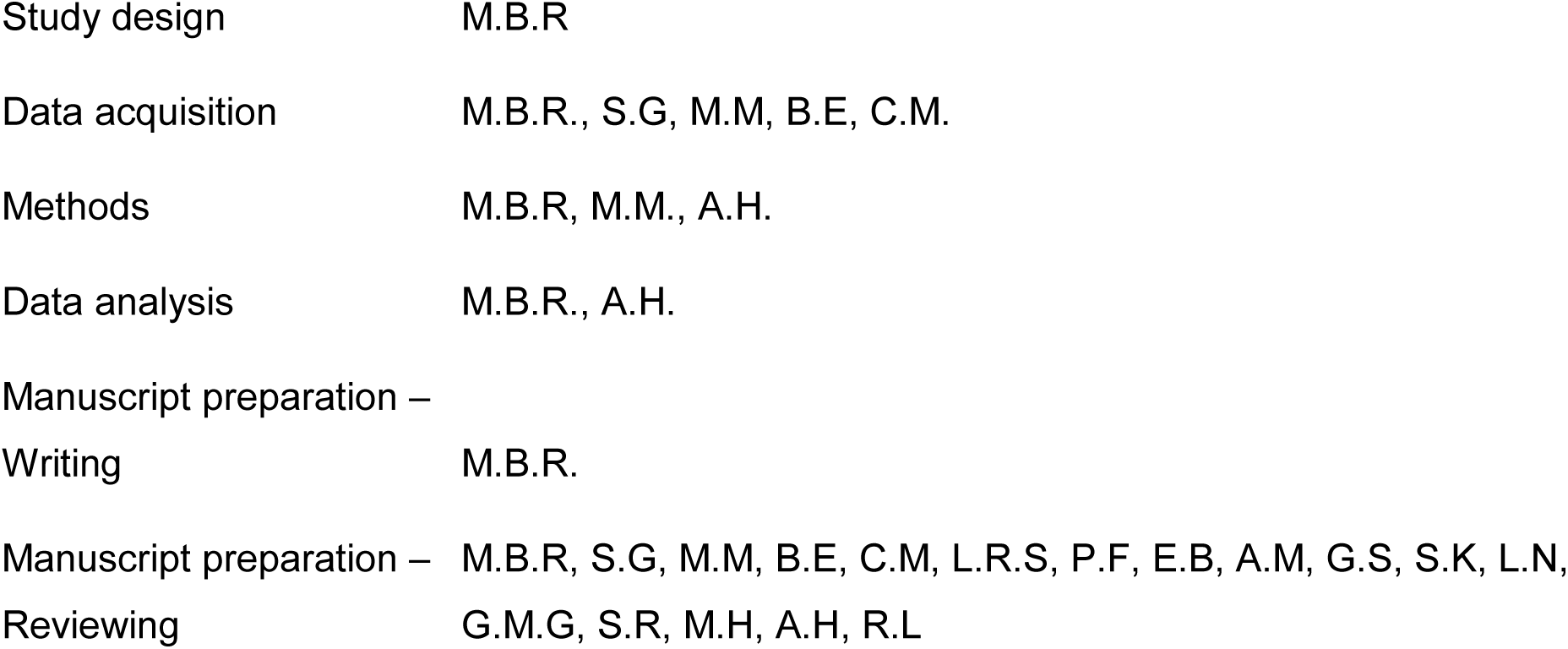

All authors discussed the implications of the findings and approved the final version of the manuscript.

### Ethics approval

Both studies were approved by the ethics committee of the Medical University of Vienna (EK 1307/2014 and 1642/2022) and conducted in accordance with the Declaration of Helsinki.

### Consent to participate

At the screening visit, all participants provided written informed consent after detailed explanation of the study protocol.

### Consent to publish

Not applicable.

### Data Availability Statement

Raw data will not be made publicly available due to reasons of data protection. Processed data and custom code can be obtained from the corresponding author with a data-sharing agreement, approved by the departments of legal affairs and data clearing of the Medical University of Vienna.

