## Supplementary Figure 1 for "High-temporal resolution metabolic connectivity resolved by component-based noise correction"

***Supplementary Material***

**Running Title: Component-based Metabolic Connectivity Estimation**

**# Correspondence to:**

Assoc.Prof. PD. Dr. Andreas Hahn, MSc

ORCID: https://orcid.org/0000-0001-9727-7580

Medical University of Vienna, Department of Psychiatry and Psychotherapy, Austria

### Supplementary Material

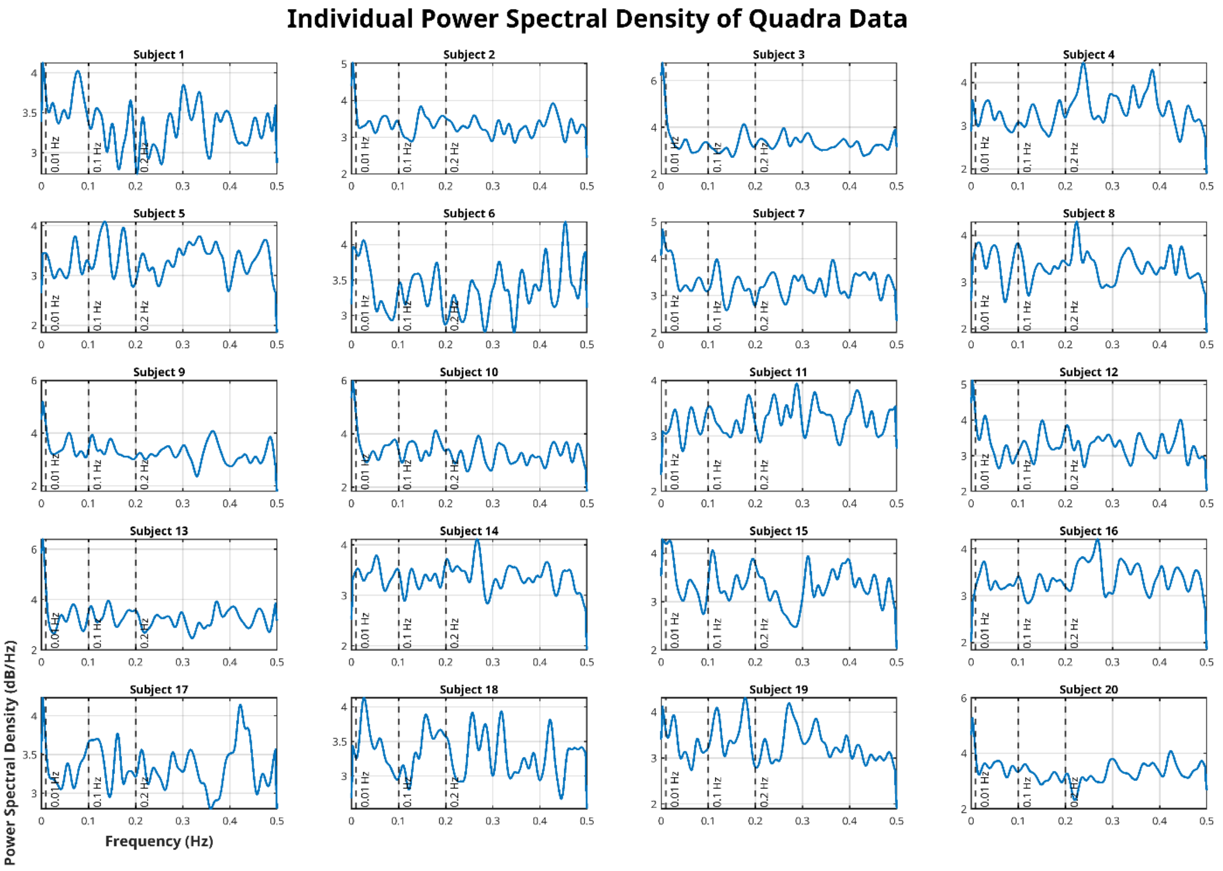


Supplemetary Figure S1: Overview of all particpants' power spectra estimates after removal of baseline uptake. Standard frequency bands are depicted as dashed vertical lines for 0.01, 0.1 and 0.2 Hz.

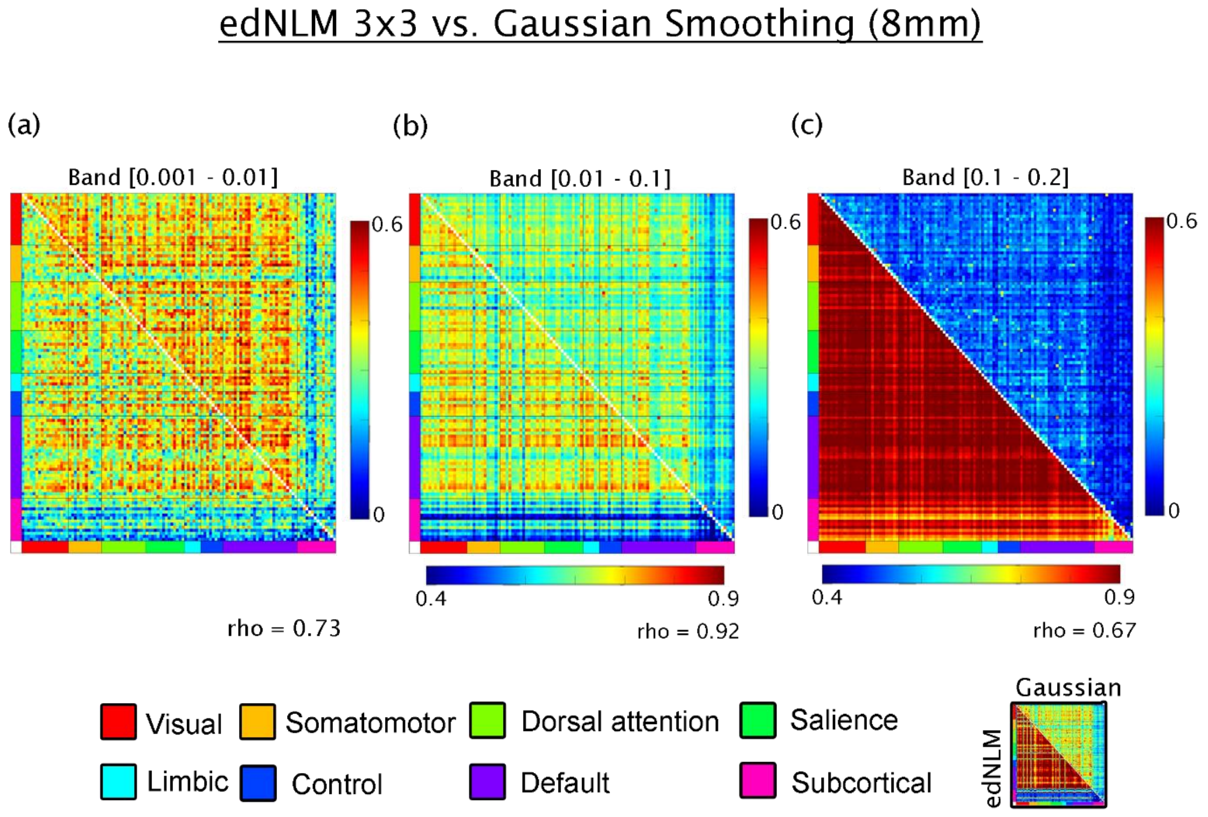


Supplementary Figure S2: Comparison of signal processing techniques applied to high-temporal resolution metabolic connectivity data acquired with a current-generation PET/MRI scanner. (a) In the lowest frequency band (0.001 - 0.01 Hz), metabolic connectivity estimates processed with extended dynamic non-local means (edNLM) filtering (upper triangle) show comparable magnitude to those processed with conventional Gaussian smoothing (lower triangle), with a moderate linear correlation between methods (Spearman's rho = 0.73). (b) In the 0.01 - 0.1 Hz band, edNLM filtering yields noticeably higher connectivity values compared to Gaussian smoothing, although the correspondence between methods remains high (rho = 0.92). (c) In the higher frequency band (0.1 - 0.2 Hz), the linear association remains moderate (rho = 0.67), but the edNLM-processed data consistently overestimate connectivity relative to the Gaussian approach. Connectivity values are grouped according to large-scale functional networks defined by [52], along with subcortical regions from the Harvard-Oxford atlas [53].
